# State-of-the-RNArt: benchmarking current methods for RNA 3D structure prediction

**DOI:** 10.1101/2023.12.22.573067

**Authors:** Clément Bernard, Guillaume Postic, Sahar Ghannay, Fariza Tahi

## Abstract

RNAs are essential molecules involved in numerous biological functions. Understanding RNA functions requires the knowledge of their 3D structures. Computational methods have been developed for over two decades to predict the 3D conformations from RNA sequences. These computational methods have been widely used and are usually categorised as either *ab initio* or template-based. The performances remain to be improved. Recently, the rise of deep learning has changed the sight of novel approaches. Deep learning methods are promising, but their adaptation to RNA 3D structure prediction remains difficult. In this paper, we give a brief review of the *ab initio*, template-based and novel deep learning approaches. We highlight the different available tools and provide a benchmark on nine methods using the RNA-Puzzles dataset. We provide an online dashboard that shows the predictions made by benchmarked methods, freely available on the EvryRNA platform: https://evryrna.ibisc.univ-evry.fr/evryrna/state_of_the_rnart/

## Introduction

Ribonucleic acids (RNAs) are macromolecules that play diverse biological roles in living organisms. RNAs are involved in numerous physiological processes, such as protein synthesis, RNA splicing, or transcription regulation, as well as in various human diseases. RNAs also have the potential to be used as therapeutic agents for different purposes, like cancer (1). Understanding RNA functions is a challenging task that has been studied for decades.

The biological function of RNA is, like protein, determined by the 3D conformation of the molecule. This folding can be determined by experimental methods like X-ray crystallography, NMR or, more recently, cryo-EM (2). Nonetheless, these methods are costly both in time and resources. On the other hand, sequencing methods (like next-generation sequencing (3)) have progressed, and a large number of sequences has become available, without any structural data. As a result, there is a huge gap between the known RNA sequences compared to the solved 3D structures. Up to December 2023, there are 7,296 solved RNA structures in the PDB (4) compared to 2,924,924 RNA sequences in Rfam (5). Only 136 out of 4,170 RNA families have at least one known structure. Therefore, computational methods have been developed for the past decades to compute RNA 3D structure from the sequence. Two main approaches have emerged: the *ab initio* and the template-based. While the first uses molecular dynamics and force fields, the latter relies on a database of known structures. None of these approaches predicts RNA structure perfectly and methods still emerge.

Since its first appearance in CASP13 (6), Alphafold (7, 8) from DeepMind has successfully predicted an enormous number of protein 3D structures. The team used deep learning techniques to predict the atomic positions of each amino acid of the sequence with high precision. Nonetheless, it can not be applied directly to RNAs due to the protein and RNA intrinsic physical differences. Indeed, the sequences are different between RNA and proteins in terms of individual elements (amino-acid compared to nucleotides), diversity of sequence range (RNA sequences range in length from a few tens to several tens of thousands of nucleotides, while proteins are a few hundred amino acids long), the number of available structure data and the stability of the folding (a given sequence of protein can fold into one stable conformation compared to multiple conformations for RNA). As a direct utilisation of AlphaFold for RNAs is not possible, works have emerged to adapt AlphaFold’s success to RNAs. The breakthrough success of AlphaFold is not yet found for RNAs (9), but some inspired works have promising performances.

Works have been done to review the state-of-the-art existing methods. A recent study (10) describes up-to-date models while highlighting the need to use probing data. Another review (11) also describes past methods and points out the detailed types of inputs that can be integrated into developed models. On the other hand, a review (12) describes only the *ab initio* methods with the force fields used for each method. A final recent review (13) discusses recent advances in terms of RNA but is not specific to the 3D structures. It sheds light on the machine learning advancements in the RNA field.

In this paper, we aim to give the reader a comprehensive overview of the RNA 3D structure prediction. Through a detailed description of *ab initio*, template-based and deep learning approaches, we detail the available tools and benchmark them on a dataset to compare their performances. The results are easily reproducible and an interface with the predicted 3D structures is provided and freely available on the EvryRNA platform: https://evryrna.ibisc.univevry.fr/evryrna/state_of_the_rnart/. The user can interact with the dashboard to select the RNA to visualize and look at the different predictions computed for the benchmark.

The paper is organised as follows: we first provide an overview of the main predictive methods developed through decades for predicting RNA 3D structure. We give a broad overview of the field and include state-of-the-art deep learning approaches, with published or preprint works. Finally, we benchmark the models available on a common dataset to assess their global performances.

## Methods

Computational methods aim to predict the atomistic positions and interactions in the RNA molecule. These methods try to reproduce the complexity of RNA, which can be single or multi-stranded (association of different strands of RNA), or even circular (where 3’ and 5’ ends are covalently linked). Computational methods tend to follow the same steps: sampling the conformational space (creation of a set of candidate structures) and discrimination of the candidates. The final structure is usually chosen with either the lowest energy or the center of a cluster of lowest energy structures. Methods can be classified as *ab initio*, template-based or deep learning-based. *Ab initio* methods integrate the physics of the system, while template-based methods are based on constructing a mapping between sequences to known motifs. Deep learning approaches use data to feed a neural network architecture that predicts RNA 3D structures from sequence or MSA (Multiple Sequence Alignment).

We present in the following a description of the state-of-the-art methods for RNA 3D structure prediction. The methods are organised by approach type (*ab initio*, template-based and deep learning) and chronologically. A timeline of all the methods, including the required inputs, is shown in Figure 1.

**Figure 1.**
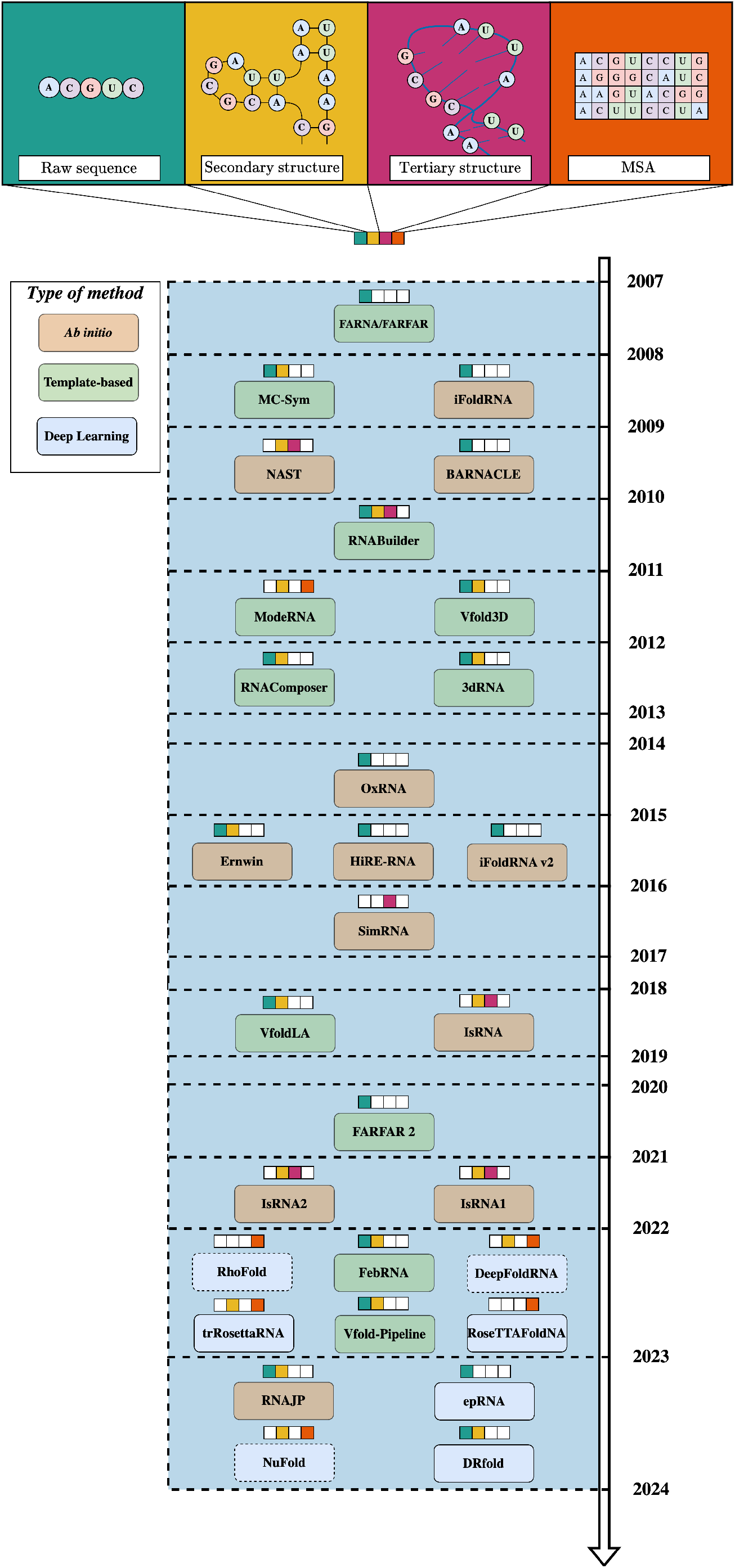
Overview of the state-of-the-art methods for predicting RNA 3D structures. The different inputs are either raw sequence, secondary structure, tertiary structure or multiple sequence alignment (MSA). Dashed methods are preprint works.

### *Ab initio* methods

*Ab initio* (or prediction-based) methods tend to simulate the physics of the system. They also capture the folding dynamics, such as energy landscapes. RNA molecules are represented at the atom level, and forces are applied to simulate real environment conditions. To explore the conformation space, sampling algorithms are used, like Monte Carlo (MC) (14) or molecular dynamics sampling (15). As the simulation can be time-consuming, a key parameter of *ab initio* methods is the granularity of the nucleotide representation. It is characterized by the number of beads per nucleotide, wherein atoms are omitted and substituted with new representative atoms. A bead refers to the number of atoms per nucleotide, which defines the granularity of the method. NAST (16), for instance, uses one atom per nucleotide, while other methods like iFoldRNA (17), OxRNA (18), HIRE-RNA (19), SimRNA (20), IsRNA1 (21), IsRNA2 (22) and RNAJP (23) tend to have more atoms per nucleotide. Other methods use different granularity like Ernwin (24) with helix as a base or BARNACLE (25) with a bayesian model.

**iFoldRNA** (17) is a three-bead per nucleotide method with discrete molecular dynamics to simulate the RNA folding process. Another version of iFoldRNA, called iFoldRNA v2 (26), adds clustering on root mean square deviation (RMSD) after simulation to reconstruct the center of founded clusters. Each bead represents a phosphate, sugar or nucleobase. The force field incorporates angle interactions, base pairing, base stacking, or hydrophobic interactions.

A web server is provided, but not the source code. The web server requires having an account. When connected, a user can make predictions from a sequence and, optionally, a 2D structure. The computation time is high: a sequence with less than 100 nucleotides takes more than one day to be processed.

**NAST** (16) models at the one-point-per-residue resolution but considers the geometrical constraints from ribosome structures before discriminating the obtained structures with root-mean-square deviation. It utilizes knowledge-based statistical potential to guide the simulation and cluster-generated structures. The bead is located at the *C*3^*′*^ atom.

No web server is provided; the source code is available and written in Python 2.

**BARNACLE** (25) is based on a Bayesian parametrized model using the seven angles characterizing a nucleotide with a hidden Markov chain process. It models marginal distributions for the dihedral angles using a mixture of probability distributions. It links the dependencies between angles with a Markov chain of hidden states. It helps reduce input representation while capturing the length distribution of helical regions.

No web server is provided, but the source code is available. We tried to run the code, but we got errors. We also tried to convert the Python 2 code to Python 3 without success.

**OxRNA** (18) is a 5-bead coarse-grained approach that uses both virtual move Monte Carlo (VCMC) and umbrella sampling (27) to sample the conformational space. It manages to characterize the thermodynamics of RNA molecules. The potential energy of the model splits terms that are nonnearest-neighbour pairs of nucleotide and neighbours. It also incorporates temperature dependence, as the coarse-grained interaction is assumed to be free energy rather than potential energy.

A web server and source code are available. Nonetheless, the source code details the web server. The required inputs for the local or web servers are of a specific format, with configuration and topology files. Therefore, it is not straight-forward to properly convert a sequence to server inputs.

**Ernwin** (24) uses Markov chain Monte Carlo (MCMC) with a helix-based model that maps the helices to cylinders and loops to close edges connected to a helix. The force field uses five energy terms like steric clash energy or knowledge-based potential of mean force.

A web server and a source code are available. The web server only returns coarse-grained molecules. There is still, up-to-date, no full-atom reconstruction included.

**HiRE-RNA** (19) shows that noncanonical and multiple base interactions are necessary to capture the full physical behaviour of complex RNAs, with a six-bead nucleotide method. It uses a model with geometric parameters determined from 200 structures. The potential integrates stacking and base-pairing terms that consider base orientations. The Replica-Exchange Molecular Dynamics (REMD) simulations are used for sample strategies.

There is no web server nor source code available.

**SimRNA** (20) uses Monte Carlo steps with a five-bead nucleotide approach guided by an energy that considers local and non-local terms. The local term includes bond length or angle interactions, while non-local terms consider base-to-backbone interactions. The sampling procedure is the asymmetric Metropolis algorithm (28). The predicted structures are based on clustering methods of lower energies.

Web server and standalone server are available. The code is well-documented and can be used easily. The web server is user-friendly, and numerous customisations can be added to the simulation. The code can be used locally but requires a lot of resources (and CPU) to be run efficiently.

**IsRNA** (29), **IsRNA1** (21) and **IsRNA2** (22) are based on a coarse-grained method with five-bead per nucleotide to predict noncanonical base pairs. The energy used includes bond length, bond angle bending and torsion angle energies. The energy also combines covalent energy functions for basepairing interactions. Non-local terms like base-base, base-backbone and backbone-backbone interactions are also included. In the IsRNA1 model, the canonical base-pairing adds interaction distances to consider bond strength compared to IsRNA. IsRNA2 better integrates noncanonical base pairing interactions in large RNAs compared to IsRNA1.

A web server is available for IsRNA1, while the source code can only be downloaded with an account. The installation requires multiple libraries that also require having an account on other websites. The web server takes multiple hours to predict hundreds of nucleotides. No web server is available for IsRNA2, and the web server for IsRNA1 starts its simulation process with structures predicted from IsRNA.

**RNAJP** (23) uses a coarse-grained approach at both atom and helix levels. It represents a nucleotide with five beads to describe the Watson-Crick, Hoogsten and sugar edges in bases. The force field used is a sum of 12 energy terms considering bonded interactions in length, bond and torsion angles, as well as base pairing and base stacking interactions. The energy integrated uses terms for the manipulation of helices and loops.

No web server is available and the source code can only be downloaded with an account. We had errors with the *bp_stk_paras* folder, where capitalization variations were missing. We managed to get the program running by modifying this folder.

Using physics-based modelling, coarse-grained approaches can predict RNA tertiary structures from raw sequences. The energy-based scoring function helps discriminate or guide predicted structures. Final predictions are usually either the lowest energy molecules or centroid of clusters. Current coarse-grained approaches fail to consider the formation of non-canonical pairs and, even more, the base side of interactions. The size of the considered RNA limits those methods: the longer the sequence, the more time-consuming the simulation is. The simulation time is not linear with the sequence length: an increase in the sequence length would highly increase the number of conformational states. Having an efficient sampling method is a challenging task and the key to efficient *ab initio* methods. The final limitation of those methods is the discriminator function, which is usually energy-based. An inaccurate energy function could result in a non-native predicted structure and bias the sampling method, which often guides the sampling procedure.

### Template-based methods

Template-based (or fragment-assembly) approaches rely on the fact that molecules that have evolution similitude adopt similar structures. A template molecule can be used as a structural basis, where other mutated sequences tend to retain similar and global conformations. A database of known RNA structures is used as a reference. Those structures have a mapping between their sequence and motif/structure/fragment. The size of the fragments considered is a key parameter for the efficiency and accuracy of the method. It can be at the nucleotide level or at the secondary structure elements (SSEs) level, for instance. Methods like RNABuilder (30) and ModeRNA (31) use one nucleotide per fragment, while FARNA/FARFAR (32) and FARFAR 2 (33) use three nucleotides per fragment. MC-Sym (34), RNAComposer (35), Vfold (36), VfoldLA (37), 3dRNA (38**–** Vfold Pipeline (42) and FebRNA (43) consider as base representation SSEs. The predicted structure can be refined to prevent clashes with energy minimization.

**FARNA/FARFAR** (32) is one of the first template-based methods to predict RNA 3D structures. It is inspired by Rosetta low-resolution protein structure prediction method (44). It uses an energy function of six terms relying on physics-based constraints, a metropolis criterion for fragment assembly using torsion angles replaced at each Monte Carlo step. While energy is computed atomistically with FARFAR, FARNA uses a simplified coarse-grained potential. Both energies can form non-canonical pairs but are limited by size and cannot predict large molecules. FARNA/FARFAR uses short segments as blocks (three-nucleotide segments) and thus needs numerous MC samplings to find a stable structure. FARFAR 2 (33) was proposed to increase the accuracy and speed. It also adds a clustering method to discriminate the most common structures.

There is a web server for FARFAR and FARFAR 2, but no source code is available. The prediction time is quite high, with multiple days for a single prediction.

**MC-Sym** (34) uses the SSEs, with nucleotide cycle modulus as blocks. It takes as inputs both raw sequence and 2D structures from MC-Fold (34) method to minimize the physics-based force field. It relies on a representation of nucleotide relationships named nucleotide cyclic motif (NCM), incorporating more context-dependent information. This representation is used to infer a scoring function for both secondary and tertiary structure prediction. A database with lone-pair loops and double-stranded NCMs is used in the pipeline and in the scoring function.

The source code is unavailable, but a well-documented web server is provided. The web server is user-friendly, and there is almost no waiting time for a job to run. However, it requires secondary structures from MC-Fold to predict 3D structures.

**RNABuilder** (30) uses multi-resolution modelling (MRM) and multibody dynamics simulation. It is based on a target-template alignment that assigns correspondences between residues and spatial constraints. It is described to predict *Azoarcus* group I intron and can be extended to other structured RNAs. It combines secondary and tertiary base pairing contacts in the force field. It can also solve structures with small connecting regions without a template.

No web server is available but a source code is available, well-documented and usable.

**ModeRNA** (31) searches for fragments in a database to replace the mutated structure before using energy minimization to refine the final structure. It uses atomic coordinates of the template and prevents backbone discontinuities by adding short fragments of other structures. It provides different strategies to build RNA structures that can be modified easily.

A web server and a code are provided. Both of them require a 3D structure as input.

**Vfold3D** (45) constructs 3D structures from fragment databases. It uses the lowest free energy secondary structures converted to known fragments. The reconstruction of fragments is coarse-grained before being converted to allatom. The final refinement of the structures uses AMBER energy minimization (46, 47). **VfoldLA** (37) uses a template database with single-stranded loops or junctions. Instead of searching for whole motifs, its granularity is finer and allows smaller blocks to be integrated. It helps prevent the limit of Vfold3D, which uses whole motifs (instead of smaller blocks) limited by the number of available RNA data. Integration of two previous methods has been done in **Vfold-Pipeline** (42). Given a sequence in input, the pipeline uses Vfold2D (48) to predict the secondary structure and then uses either Vfold3D or VfoldLA for the final 3D structure prediction.

A web server is available for either Vfold3D, VfoldLA and Vfold-Pipeline. The source code is also available and usable.

**RNAComposer** (35) creates a database (named RNA FRABASE (49)) with fragment mapping 2D elements to 3D motifs before using refinement. The SSEs are used as minimum blocks to assemble the different fragments. The method uses the Kabsch algorithm (50) to assemble the 3D structure elements. The refinement of the structure concatenates two energy minimization methods: torsion angles energy (using CYANA (51)) and atom coordinate with CHARMM (52).

There is a web server accessible, but no source code is provided.

**3dRNA** (38) uses a fragment assembly approach guided by a scoring function, 3dRNAScore (53), where the SSEs considered are improved by more base pairs from connected stems. It uses SSEs as blocks and predicted structures with a clustering approach using 3dRNAScore as criteria. Improvements have been made over the years (39–41) with, for instance, an increase of about ten times the number of templates in the 3D template library (41). It also adds the possibility to predict circular RNAs.

A web server is provided, and the source code is available only after login. It is required to have other software installed to run the standalone code.

**FebRNA** (43) creates a 3D fragment ensemble and identifies the 3D coarse-grained structure using cgRNASP (54) score, with three-bead per nucleotide. It performs all-atom reconstruction followed by refinement. The building of fragments is executed with secondary structure tree (SST) (55), where each stem is considered as a node of a tree structure. A 3D structure is build through sequential superposition between coarse-grained atoms of a loop and stem according to the SST order.

No web server is accessible, but the source code is available and well-documented. Nevertheless, we did not manage to run the code because we had errors.

Template-based methods allow the prediction of RNA 3D structures with the help of available data. They create a database mapping sequence to fragments (or motifs) before assembling it to refine final structures. However, the number of experimental RNA structures is a bottleneck for the good accuracy of the models. Templates like SSEs tend to be inaccurate or missing in the constituted database, preventing good predictions of structures. They also fail to generalize to unseen structures. As many RNA families have not yet been discovered, such approaches would probably fail to predict new families.

### Deep learning approaches

In the CASP competition, an end-to-end approach has been introduced and overperformed all previous works for predicting protein 3D structure: AlphaFold (7, 8). It has changed the structural biology field and raised the interest of researchers. Recent works have been done to predict RNA 2D structures (56–58), as the available data is much higher than solved 3D structures. Other deep learning works try to predict energy function (59, 60), while others infer torsion angles from the sequence (61). Such angles can nevertheless be used to help the prediction of 3D structures. Preprint works have been released like DeepFoldRNA (62), RhoFold (63), and NuFold (64) to predict 3D structures with attention-based (65) methods. Four deep learning approaches, epRNA (66), DRfold (67), RoseTTAFoldNA (68) and trRosettaRNA (69), have recently been published. As advancements in the field are moving fast, we describe both preprint and published works in the following.

**DeepFoldRNA** (62) is a preprint work that predicts RNA structures from sequence alone by coupling deep self-attention neural networks with gradient-based folding simulations. It predicts distance and orientation maps, as well as torsion angles, with transformer-like blocks. It uses MSA and 2D structure as inputs. A BERT-like (70) loss was also implemented to make the model more robust. A self-distillation approach is used to get around the lack of data. It incorporates bp-RNA-1m (71) sequences to predict their structures and integrate them into the training set. To convert the neural network outputs to 3D structures, they use L-BFGS (72) folding simulations with energy defined by the weighted sum of the negative log-likelihood of the binned probability predictions.

A web server and a source code are provided. We tried to predict sequences from the web server but never received the results.

**RhoFold** (63) is a preprint work with an end-to-end differentiable approach for predicting RNA 3D structures. The model’s input is the MSA, and features are extracted with a pre-trained model RNA-FM (73) (trained over more than 23 million sequences). RNA-FM gives an MSA coevolution matrix and pairwise residue features. A module called E2EFormer with gated attention layers is applied to predict the main frame 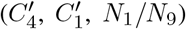 in the backbone and four torsion angles (*α, β, γ, ω*). An IPA (invariant point attention) is used in modelling 3D positions. It predicts each frame’s rotation and translation matrices based on the sequence and pair representation from the E2Eformer module. Given the predicted frames and angles, the structure module can generate the full-atom coordinates of an RNA without simulation. It also uses self-distillation with bp-RNA-1m (71) and combines the training process with a loss that takes into account 1D (sequence masking), 2D and 3D (Frame Aligned Point Error (FAPE)) elements.

A web server and a source code are provided. The web server is easily usable, while the standalone code requires more than 500 GB of space to download the database, even for inference.

**RoseTTAFoldNA** (68) is a published work with an end-to-end deep learning approach that predicts 3D structure for RNA molecules and protein-DNA and protein-RNA complexes. It incorporates three representations of molecules: sequence (1D) with MSA representation, residue-pair distances (2D) and cartesian coordinates (3D). The 3D representation uses the position and orientation of phosphate, as well as torsion angles. The model can take as input protein, DNA and RNA. It was trained on five types of structures: protein structures, AlphaFold2 predictions, protein complexes, protein/NA complexes and RNA structures. The network was first trained before being fine-tuned, where energy terms were added to the loss of the network.

A source code is provided, but no web server exists. The source code requires more than 500Gb of free space to download sequence and structure databases.

**trRosettaRNA** (69) is a published work inspired by two methods for 3D protein structure prediction, AlphaFold2 (8) and trRosetta (74– It uses MSA and secondary structure (predicted by SPOT-RNA (77)) as inputs. The network architecture is inspired by AlphaFold2 Evoformer block and thus uses transformer networks. The full atom reconstruction uses energy minimization with restraints from predicted geometries weighted by parameters optimized from random RNA from the training set. The model is trained on PDB data with sequences that have homologs. It uses bpRNA (71) from Rfam (5) for self-distillation to increase the available data. Distillation is regulated with a Kullback-Leibler divergence.

A web server is available, but no standalone code.

**epRNA** (66) is a published work with an Euclidean parametrization-based neural network that predicts RNA tertiary structure from sequence only. It is trained to predict a distance matrix that is then added to the loss. The network uses convolutional networks and uses one hot encoding as input. epRNA uses RNAs from the PDB and splits them into training and test sets (60% for training and 40% for testing). The method achieves invariance in terms of rotation and translation, but not for the reflection of the molecule. It means that the mirror image of a chiral molecule is chemically distinct, but this distinction is not made in the network.

A source code is available, but no web server. The code is easy to use, and the installation process is straightforward. There is no need to install huge datasets to perform predictions.

**NuFold** (64) is a preprint work with an adaptation of AlphaFold2 work for RNAs. It considers the base frame with four atoms: *O*4^*′*^, *C*1^*′*^, *C*4^*′*^ and either *N* 1 (for C and U) or *N* 9 (for G and A). It also adds heads to predict the distance between *C*4^*′*^ and *P* atoms, and the torsional angles to help the full-atom reconstruction. It uses as inputs MSA and secondary structure predicted by IPknot (78). The NuFold network comprises two key components: the EvoFormer block and the structure model. The EvoFormer part is a transformer model that embeds information into single and pair representations. The structure model converts the embedding into 3D structures. It is recycled three times to increase the accuracy of predictions. The network outputs are the translation and rotation of the four base frames and torsion angles. The torsion angles guide the reconstruction of full-atom representation from the base frames.

No web server is available, and no code yet. It is said that the code will be available after a clean-up by the authors.

**DRfold** (67) is a published work with an end-to-end transformer-based approach that takes as input RNA sequence and secondary structure. It uses a three-bead representation for a nucleotide. It converts the inputs into sequence and pair representations before feeding them to transformer blocks. A structure module outputs frames converted to FAPE (frame aligned point error) potential, while a geometry module predicts rotation and translation property converted to geometry potentials. These predicted frame vectors and geometry restraints are aggregated to a potential for structure reconstruction. The final step includes all-atom reconstruction and refinement using Arena (79) and OpenMM (80).

No web server is provided, but a source code is available. It requires the download of numerous libraries.

Deep learning methods are promising and have good performances on testing datasets. Nonetheless, deep learning models need a huge amount of data, which is unavailable for RNA 3D structures. To avoid this bottleneck, methods use self-distillation. They also mainly input MSA representation like AlphaFold. MSA remains a limitation as the number of known RNA families is restricted. The overall quality of the predicted structures remains to be validated with new data from unseen families.

A summary of the state-of-the-art tools, including information on their implementation, is given in Table 2. We have added a column mentioning whether the methods explicitly predict multi-stranded RNAs. Only one method explicitly points out the fact that they predict circular RNAs: 3dRNA.

## Results

In this section, we detail the results of available methods for RNA 3D structure prediction. To have a fair comparison between existing methods, we benchmark them on three different test sets. We evaluated and compared the predicted structures using standard metrics described in a previous work (81).

### Benchmarked tools

As summarized in Table 2, some of the state-of-the-art methods do not have a web server or a standalone code available. It is the case of Hire-RNA (19) and NuFold (64). Among the remaining tools, unfortunately, many are hard to use or not working. Among the available standalone codes, we only manage to run RNAJP (23). DeepFoldRNA (62), FebRNA (43) or RoseTTFoldNA (68) require the download of databases. Those databases could have more than 500Gb and thus be hardly usable for users. Ernwin (24) and epRNA (66) only return coarse-grained structures and thus increase the use complexity. Among the web servers available, ModeRNA (31) needs as input an initial 3D structure, which we did not have for the benchmark (and would also bias the comparison with the other methods). OxRNA (18) requires a specific input format, which makes it hard for the user to use. FARFAR 2 (33) has a web server with a computation time too long to be included (multiple days of predictions), where our predictions did not lead to results. DeepFoldRNA (62) and Drfold (67) have web servers where we did not get the structures after making the request. The server of iFoldRNA (17) is very hard to connect to and failed to perform all the predictions: we were only able to have a few predictions As a benchmark, we thus considered the remaining ten methods described in Table 2. We used RNA-tools (82) to clean the predicted structures and to normalize them. This software enables the operation of RNA structures and allows their standardisation to help better evaluate them. All methods were used with their web servers except for RNAJP, which was used locally. We set a computation limit for RNAJP computation (50 *×* 10^6^ steps in the simulation). For SimRNA, we stop the simulation at 20000 frames. We used the web server for Rhofold, which does not use MSA and might get lower performances than the MSA version.

Not all tools could predict directly from the sequences, a secondary structure is required. We decided, when needed, to use the secondary structure predicted by MXFold2 (83), a recent deep learning-based tool giving good prediction results. The choice of MXFold2 was arbitrary but should be consistent between the models to have a fair comparison. For MC-Sym, it is required a secondary structure from MC-Fold (34).

### Test Sets

To compare the models’ performances, we used three different test sets. We considered single-stranded RNAs to enable the comparison between all the models. The aim of the benchmark is to enable the comparison of user-available models, where no specific parameters optimization is performed for the predictions. We kept RNA with sequence below 200 nucleotides, except for two RNAs from the first test set (sequence length of 210 and 298 nucleotides), to have a complex enough dataset for the comparison.

The first test set, which we call Test Set I, is a non-redundant dataset of RNA structures from RNAsolo (84). We downloaded the representative RNA molecules from RNA-solo with a resolution below 4Å and removed the structures with a sequence identity higher than 80%. Then, we considered only the structures with a unique Rfam family ID (5), leading to 29 non-redundant RNA molecules, with a sequence between 40 and 298 nucleotides. The details of each PDB ID and Rfam family from Test Set I are described in

Table S1. Nonetheless, this dataset does not ensure that there has been no data leakage in the training of the different models.

We also included predictions from a collaborative test set from the community: RNA-Puzzles (85–88), which we refer as Test Set II. The RNA molecules proposed through the years as a challenge are solved structures that have challenging properties: multi-stranded structures, ribozymes, riboswitches and more. We considered single-stranded RNAs, which represent 22 RNAs with a sequence between 27 and 188 nucleotides. As some predictions are available in the published results of RNA-Puzzles, they are results of the optimization of parameters from each group, which is nearly available for users. As this benchmark aims to report results on user-available solutions, we included predictions we made from the tools, easily reproducible for this benchmark. More details about the considered RNAs, as well as their families, are given in Table S2.

A collaboration between RNA-Puzzles and CASP teams led to the CASP 15 competition (89). 12 RNA targets were proposed. As four targets exceed 200 nucleotides, most of the models fail to predict these structures. Therefore, we considered the eight target RNAs with sequence lengths below 200 nucleotides, which we named Test Set III. This dataset aims to evaluate the robustness of the methods. Details on the structures for Test Set III are available in Table S3.

### Evaluation metrics

To evaluate and compare the quality of predictions, we used different metrics. Each metric has its specificity, which is why we computed most of the available metrics using RNAd-visor (81).

The first metric is the well-known RMSD (Root-Mean-Square-Deviation), which is very sensitive to local differences. The INF (Interaction Network Fidelity) (90) metric incorporates RNA key interactions to evaluate RNA 3D structures better. Details on the INF score are included to depict the type of interactions that are conserved in the prediction: canonical Watson-Crick interactions (INF-WC), non-canonical interactions with non-Watson-Crick base pairs (INF-NWC) and stacking in helices interactions (INF-STACK). The INF-ALL metric summarises all these interactions into one value. The *ϵ*RMSD (91) is another tentative to incorporate RNA specificities. The TM-score (92, 93) and lDDT (94) are, respectively, the normalisation of atom deviation metric and interatomic differences, both inspired by protein evaluation metrics. Other common metrics inspired by proteins are the GDT-TS (95) (accounts for superimposition with different distance cutoffs with aligned structures) and the CAD-score (96) (measures the structural similarity in a contact-area function). To compare the torsional angle deviation that characterizes RNA molecules, the MCQ (mean of circular quantities) (97) can be computed. Finally, the P-VALUE (98) assesses if a prediction is better than a random one.

RMSD, *ϵ*RMSD, MCQ, DI, and P-VALUE metrics have good results when the values are low, whereas high values are better for INF, lDDT, GDT-TS and TM-score.

### Benchmark results

We present here the prediction results obtained by each of the tools summarized in Table 1. The predictions are reported according to the different metrics presented above. We had struggled to get secondary structures for long sequences using MC-Fold (99), and we decided to exclude MC-Sym in the comparison for Test Set III (as we managed to predict only two structures).

**Table 1.**
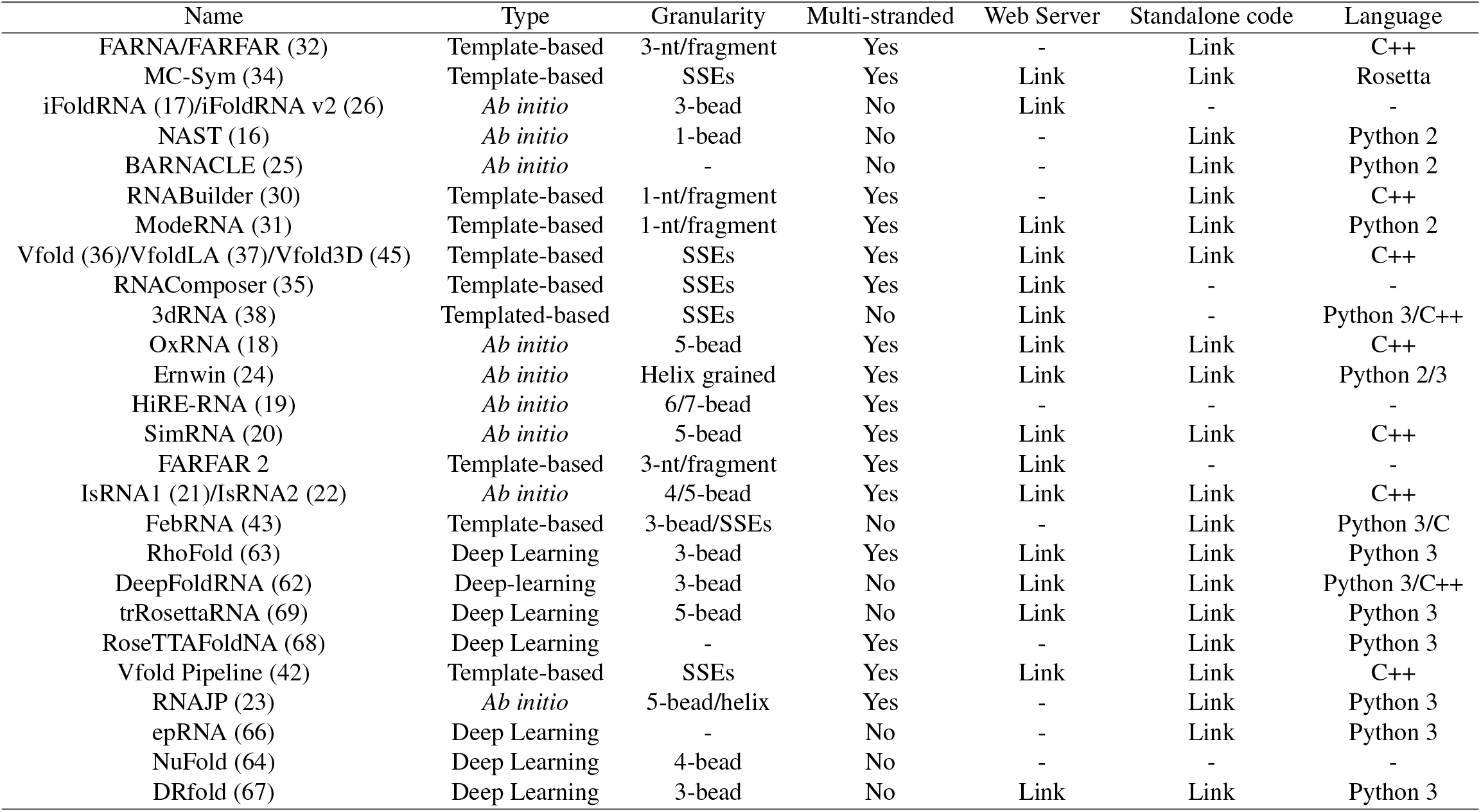
Summary of the state-of-the-art softwares for the prediction of RNA 3D structures. For each method is provided its type, granularity level, availability and implementation. We also mention if the method deals with multi-stranded RNAs.

| Name | Type | Granularity | Multi-stranded | Web Server | Standalone code | Language |
| --- | --- | --- | --- | --- | --- | --- |
| FARNA/FARFAR (32) | Template-based | 3-nt/fragment | Yes | - | Link | C++ |
| MC-Sym (34) | Template-based | SSEs | Yes | Link | Link | Rosetta |
| iFoldRNA (17)/iFoldRNA v2 (26) | *Ab initio* | 3-bead | No | Link | - | - |
| NAST (16) | *Ab initio* | 1-bead | No | - | Link | Python 2 |
| BARNACLE (25) | *Ab initio* | - | No | - | Link | Python 2 |
| RNABuilder (30) | Template-based | 1-nt/fragment | Yes | - | Link | C++ |
| ModeRNA (31) | Template-based | 1-nt/fragment | Yes | Link | Link | Python 2 |
| Vfold (36)/VfoldLA (37)/Vfold3D (45) | Template-based | SSEs | Yes | Link | Link | C++ |
| RNAComposer (35) | Template-based | SSEs | Yes | Link | - | - |
| 3dRNA (38) | Templated-based | SSEs | No | Link | - | Python 3/C++ |
| OxRNA (18) | *Ab initio* | 5-bead | Yes | Link | Link | C++ |
| Ernwin (24) | *Ab initio* | Helix grained | Yes | Link | Link | Python 2/3 |
| HiRE-RNA (19) | *Ab initio* | 6/7-bead | Yes | - | - | - |
| SimRNA (20) | *Ab initio* | 5-bead | Yes | Link | Link | C++ |
| FARFAR 2 | Template-based | 3-nt/fragment | Yes | Link | - | - |
| IsRNA1 (21)/IsRNA2 (22) | *Ab initio* | 4/5-bead | Yes | Link | Link | C++ |
| FebRNA (43) | Template-based | 3-bead/SSEs | No | - | Link | Python 3/C |
| RhoFold (63) | Deep Learning | 3-bead | Yes | Link | Link | Python 3 |
| DeepFoldRNA (62) | Deep-learning | 3-bead | No | Link | Link | Python 3/C++ |
| trRosettaRNA (69) | Deep Learning | 5-bead | No | Link | Link | Python 3 |
| RoseTTAFoldNA (68) | Deep Learning | - | Yes | - | Link | Python 3 |
| Vfold Pipeline (42) | Template-based | SSEs | Yes | Link | Link | C++ |
| RNAJP (23) | *Ab initio* | 5-bead/helix | Yes | - | Link | Python 3 |
| epRNA (66) | Deep Learning | - | No | - | Link | Python 3 |
| NuFold (64) | Deep Learning | 4-bead | No | - | - | - |
| DRfold (67) | Deep Learning | 3-bead | No | Link | Link | Python 3 |

**Table 2.** Benchmarked tools. The state-of-the-art tools are listed from the less to the most recent. Each tool is given its inputs and its method type. Seq refers to the raw sequence, and 2D for the secondary structure.

| Model | Inputs | Method Type |
| --- | --- | --- |
| MC-Sym (34) | Seq+2D | Template-based |
| Vfold3D (45) | Seq+2D | Template-based |
| RNAComposer (35) | Seq+2D | Template-based |
| SimRNA (20) | Seq+2D | *Ab initio* |
| 3dRNA (41) | Seq+2D | Template-based |
| IsRNA1 (21) | Seq+2D | *Ab initio* |
| RhoFold (63) | Seq | Deep Learning |
| trRosettaRNA (69) | Seq | Deep Learning |
| Vfold-Pipeline (42) | Seq+2D | Template-based |
| RNAJP (23) | Seq+2D | *Ab initio* |

The normalized mean of the metrics is reported for the three test sets in Figure 2, as well as for the pooled test set (all test sets gathered). We applied the min-max normalisation over the whole datasets, and reversed the decreasing metrics. Thus, each shown metric has values between 0 and 1, where 1 means best predictions and 0 is worst predictions.

**Figure 2.**
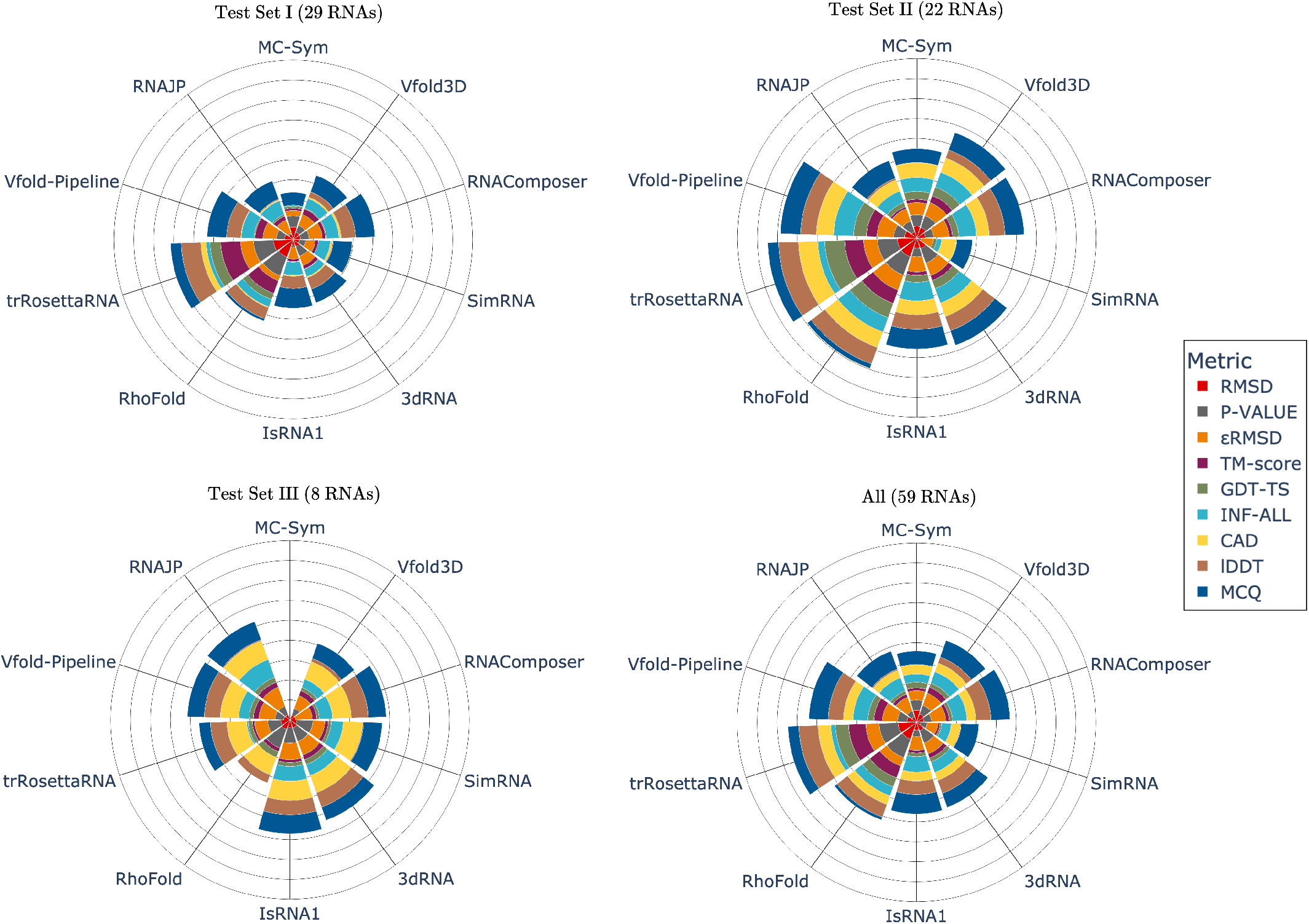
The normalised mean of metrics for each of the benchmarked methods on the different datasets. The pooled test set is named “All”. For each metric, we normalised by the min-max to ensure values are between 0 and 1, and we reverse the order for descending metrics (RMSD, *ϵ*RMSD, P-VALUE and MCQ). For a given metric, a model with a score near 1 means it has the best score compared to the other models.

trRosettaRNA outperforms the other methods in terms of cumulative metrics for Test Set I and Test Set II. It is followed by Rhofold and Vfold-Pipeline, which are almost similar for Test Set I and Test Set II. Results remain low for Test Set I compared to the two other test sets. While having good RMSD values, the deep learning approaches do not have the best INF and MCQ scores (in all test sets). It means the deep learning approaches can have a general idea of the skeleton structures, but hardly reproduce the specific key RNA interactions. It is confirmed in Table 3, where the non-Watson-Crick (non-WC) (non-canonical interactions) and stacking interactions (non-covalent interactions between adjacent nucleotide bases) are always better reproduced for *ab initio* or template-based methods (RNAComposer and IsRNA1 for non-WC, SimRNA, Vfold-Pipeline and RNAJP for stacking interactions).

**Table 3.** Metric values for INF-WC, INF-stack and INF-NWC for the different bench-marked models. Each value is given for the three test sets, separated by a “/”. We excluded MC-Sym for Test Set III, as we did not get enough predictions.

|  | INF-WC | INF-STACK | INF-NWC |
| --- | --- | --- | --- |
| MC-Sym | 0.26/0.67/- | 0.54/0.64/- | 0.14/0.06/- |
| Vfold3D | 0.60/0.67/0.55 | 0.58/0.58/0.56 | 0.00/0.00/0.00 |
| RNAComposer | 0.62/0.75/**0.73** | 0.62/0.67/0.65 | **0.21/0.33/0.10** |
| SimRNA | 0.57/0.78/0.59 | 0.65/**0.70**/0.65 | 0.08/0.07/0.10 |
| 3dRNA | 0.59/0.74/0.62 | 0.57/0.66/0.61 | 0.07/0.25/**0.12** |
| IsRNA1 | 0.61/**0.80**/0.65 | 0.64/0.68/0.65 | 0.01/0.00/0.00 |
| RhoFold | 0.50/0.70/0.31 | 0.60/0.66/0.54 | 0.10/0.20/0.03 |
| trRosettaRNA | **0.69**/0.78/0.55 | 0.57/0.63/0.55 | 0.13/0.20/0.06 |
| Vfold-Pipeline | 0.62/0.78/0.63 | 0.61/**0.70**/0.56 | 0.15/0.29/0.00 |
| RNAJP | 0.59/0.55/0.69 | **0.67**/0.65/**0.72** | 0.19/0.15/**0.12** |

For Test Set III, the deep learning models do not achieve good results, and the best method seems to be IsRNA1, followed by RNAJP, Vfold-Pipeline and 3dRNA. The worst method is RhoFold, showing difficulties in having robust predictions.

For the pooled test set, this is trRosettaRNA, which performs better overall. It is followed by Rhofold and Vfold-Pipeline. Both deep learning methods have lower values of MCQ and INF compared to Vfold-Pipeline. MC-Sym and SimRNA seem to perform worse than the other methods, which could be explained by the lack of simulation time. They still produce results with better INF and MCQ values than the deep learning approaches.

Details on the different benchmarks and results obtained are provided in the Supplementary file. Mean values for each method are described in Table S4 (Test Set I), Table S5 (Test Set II), Table S6 (Test Set III) and Table S7 (All). The associated distribution for each metric is illustrated in Figure S1 (Test Set I), Figure S2 (Test Set II) and Figure S3 (Test Set III). Detailed results of each method for each RNA are available in Figure S4 (Test Set I), Figure S5 (Test Set II) and Figure S6 (Test Set II).

To illustrate and compare visually the predictions obtained by each of the considered methods, we arbitrarily selected a structure from the RNA-Puzzles challenge: puzzle rp03, a Riboswitch (PDB ID: 3OWZ). The predicted structures, as well as the native structure, are shown in Figure 3. The detailed metrics for each prediction are available in Table S8. We did an alignment to show them on the same scale using the matching tool of Chimera (100). The model that seems to superimpose the reference structure well is trRosettaRNA, with an RMSD of 2.38. We observe good visual folding for the deep learning models and Vfold-pipeline. On the other hand, RNAJP and RNAComposer predictions do not seem to fit well with the native shape. The metric values for each model for this RNA are given in Table S8.

**Figure 3.**
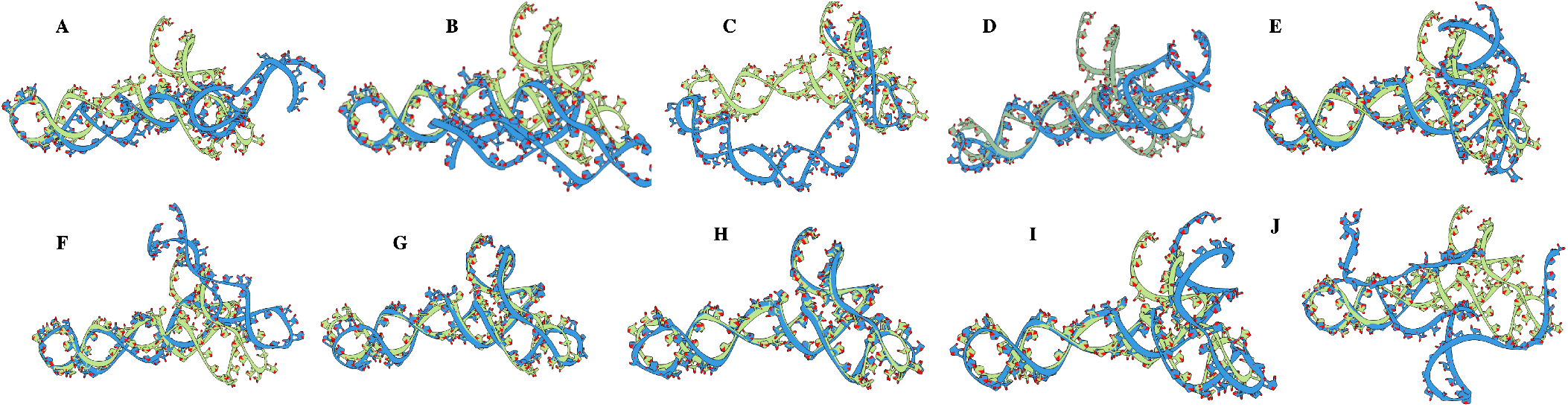
Predicted structures (in blue) for RNAPuzzle 03 (rp03) (id: 3OWZ, length: 84 nucleotides) compared to native structure (in green) using state-of-the-art methods. A: MC-Sym. B. Vfold3D. C: RNAComposer. D: SimRNA. E: 3dRNA. F: IsRNA1. G: RhoFold. H: trRosettaRNA. I: Vfold-Pipeline. J: RNAJP. Alignment was done using CHIMERA (100) and Needleman-Wunsh algorithm (101).

### Computation time

As stated above, except RNAJP, all benchmarked tools are available only as web servers. Therefore, a precise comparison of computation time performances is not possible. We thus report here for each tool the rough computation time we measure for processing a given RNA.

Figure 4 summarizes the rough inference computation time to predict RNA 3D structures for each model. We report the computation time for the RNAs with the shortest and the most extended sequences (RNA that are successfully predicted for all the methods). Vfold3D and Vfold-Pipeline have similar computation times: Vfold3D and Vfold-Pipeline are almost the same models; the only difference is the use of VfoldLA when Vfold3D does not provide predictions in Vfold-Pipeline. We observe that the *ab initio* methods have a computation time higher than the template-based and deep learning methods. This is due to the simulation processes that require a high number of computation steps. The template-based methods almost always return a structure with less than 2 hours of computation (including the queue in the web servers). On the other hand, deep learning methods tend to be very fast for inference. RhoFold predicts with high throughput, and what is the most time-consuming is the relaxation of the prediction. The *ab initio* methods are the slowest ones, with a minimum time of two hours. They often propose advanced parameters for the computation, like chemical probing restraints, distance restraints, or even freezing some residues (like those proposed in SimRNA).

**Figure 4.**
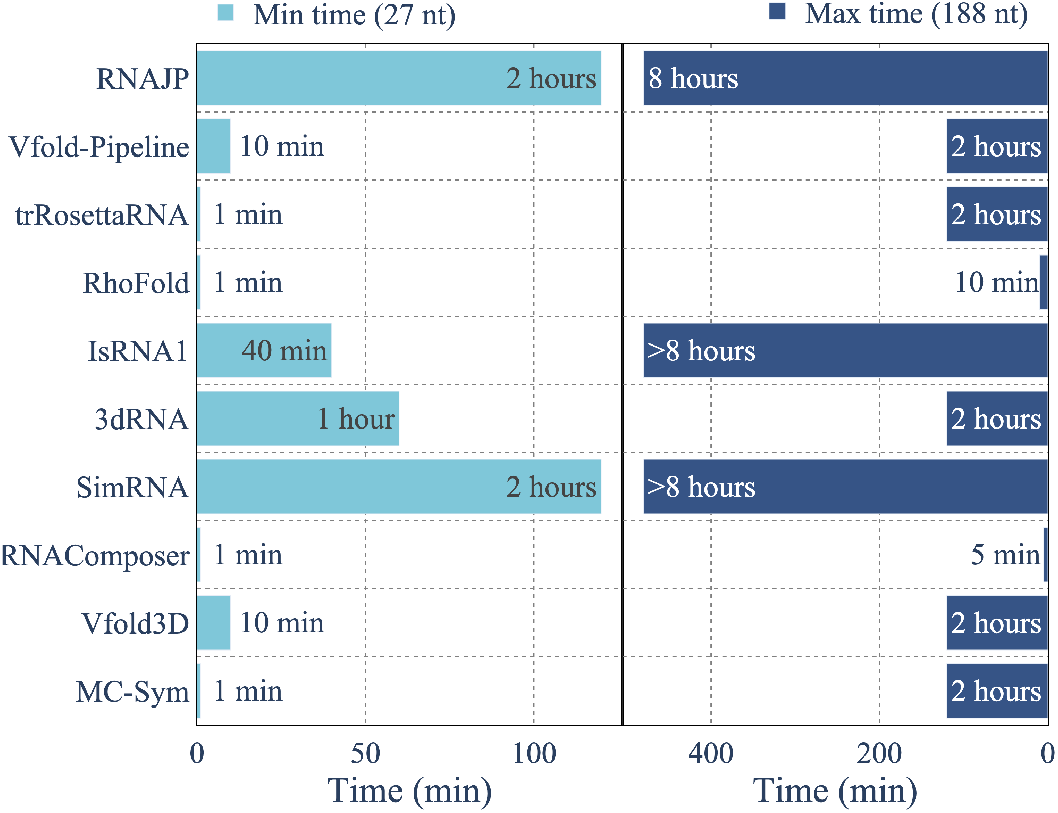
Approximate time for computation of RNA-Puzzles structures. The minimum time is for an RNA of 27 nucleotides, while the maximum time is computed for an RNA of 188 nucleotides. The computation time is an approximation, as it was run on web servers and might be slowed down by other pending jobs. The time reported for RhoFold is with the relaxation (which is slower than the raw prediction). RNAJP computation time is computed locally with a simulation time set to 50 *×* 10^6^ steps. IsRNA1 maximum time is around 15 hours, and SimRNA maximum computation time is around two days.

### State-of-the-RNArt Dashboard

We provide a dashboard (illustrated in Figure 5) with different visualisations of the predicted structures for the nine benchmarked models. The dashboard, called State-of-the-RNArt, is freely available on the EvryRNA platform: https://evryrna.ibisc.univ-evry.fr/evryrna/state_of_the_rnart/. The user can choose which RNA to compare the predictions from among the different challenges of RNA-Puzzles (85–88). We also included some of the predictions we made on the CASP-RNA (89). We make available all the obtained predictions and their evaluation with the different metrics. The State-of-the-RNArt Dashboard allows thus the reproducibily of our benchmarks and a quick visualization of the obtained 3D structures.

**Figure 5.**
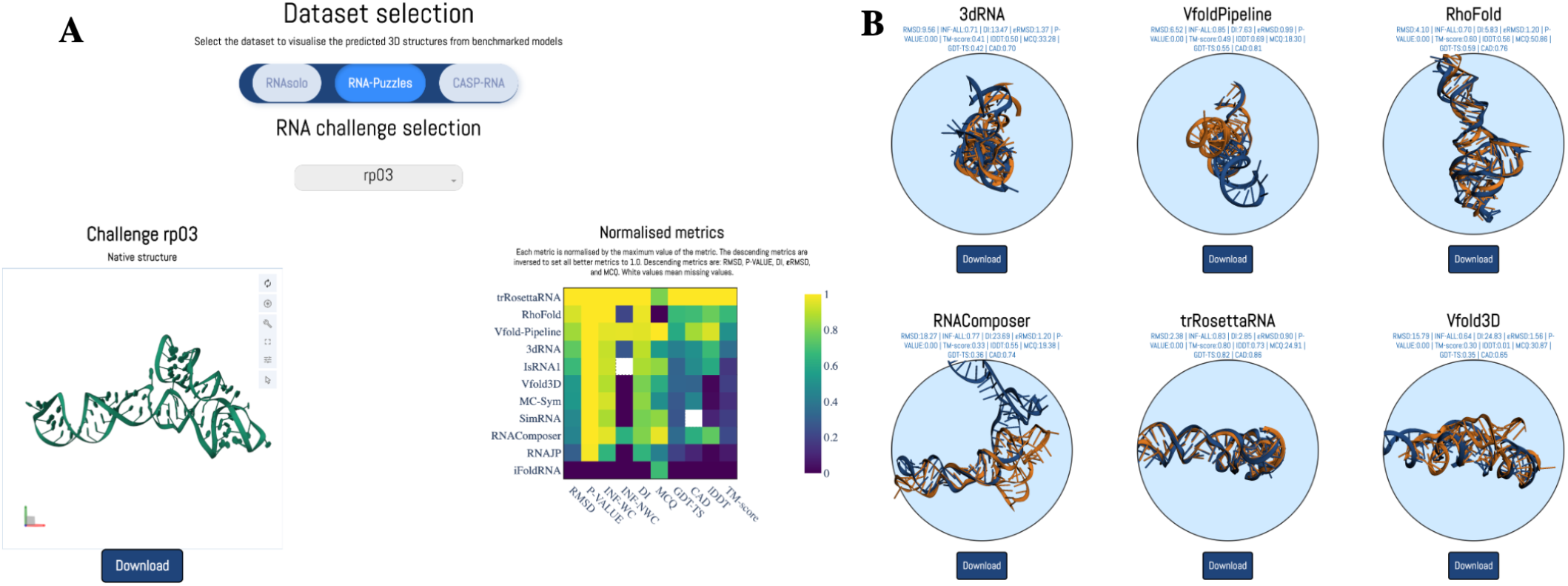
Screenshot of the State-of-the-RNArt dashboard. A: The user can choose the RNA (or challenge) with its native structure to process with the different RNA 3D structure prediction tools. It is associated with the predicted metrics normalized by the maximum value for each metric for the given RNA challenge. B: 3D visualisations of the different predictions of the benchmarked models. The native structure is coloured in orange, while the predictions are in blue. The predictions are superimposed with the native structure for visualisation using the US-align (102) tool. The associated metrics are also shown on top of the structures.

## Discussion

*Ab initio* methods are physic-based approaches that incorporate different levels of granularity in nucleotide representation. The coarse-grained approach is a trade-off between efficiency in the representation and accuracy in the prediction. We found these methods harder to use in practice as the simulation process can be very time-consuming. The standalone codes are usually unavailable or difficult to run, and good results would require high computation time and resources. They have the advantage of allowing customisation in the simulation, with the easy integration of constraints. They also predict RNA structures with more native features with better conservation of torsional angles than deep learning methods. It might be explained by the lack of data to create good guiding functions. Further development of *ab initio* models could incorporate a coarse-grained approach with efficient sample procedure and well-chosen force-field. It must be associated with full-atom reconstruction methods, adapted and efficient. Template-based methods try to map sequences to structural motifs before merging them into a whole structure, which is then refined. These methods are more efficient than the *ab initio* while still being limited. Their usage is easier than *ab initio* methods, but standalone codes remain hard to reproduce locally. Improvement of template-based methods could be based on the addition of existing physics-based methods that can predict structures not already seen. It could alleviate the prediction of unseen structures. Refining the structure after assembling could also be improved to best include fragments.

The performances of deep learning approaches seem promising. By using available data and self-distillation procedures, they perform well on the RNA-Puzzles and RNA-solo datasets. They fail, like the other two approaches, on the CASP-RNA dataset. Their performances remain incomparable to AlphaFold for proteins, and the next AlphaFold for RNA has not yet been found (9). Their usage in terms of web servers is very user-friendly: only a sequence is required, and the prediction is made very quickly. We regret the standalone codes that often require the download of a huge dataset, which is almost non-feasible for standard users. These methods are limited by a common neural network drawback: interpretability. Knowing the folding process would highly increase RNA understanding and is a step the community would appreciate. The integration of physics into deep learning methods could help reduce the black box trap as well as prevent models from overfitting.

Hybrid methods are a direction that is taken by the community with recent solutions (103) proposed in CASP-RNA (89). For instance, the second best solution from CASP-RNA (**?**) uses structures predicted by template-methods VfoldLA (37) and Vfold3D (45) before using coarse-grained simulations from IsRNA (21, 22, 29) and RNAJP (23). Hybrid solutions are usually a mix of previous methods to take the best of each of them. These recent methods are not yet available to users, so we did not include them in our benchmark.

All the previously discussed models still need to be improved with the possibility of outputting multiple structures corresponding to environment-dependent RNA molecules. Works remain to allow the prediction of long non-coding RNAs, as well as the non-canonical interactions that are still a challenge. Limitations for the classification of non-canonical base pairings can be explained by the lack of 3D data, where the systems hardly incorporate these specificities. The sequence length is still a bottleneck, where integration of all possible interactions increases the complexity and limits existing models. The predictions of multi-stranded and circular RNAs remain limited: more than half of the methods can predict multi-stranded RNAs, but only one for the circular RNAs.

## ACKNOWLEDGEMENTS

This work is supported in part by UDOPIA-ANR-20-THIA-0013 and performed using HPC resources from GENCI/IDRIS (grant AD011014250). It was also partially supported by Labex DigiCosme (project ANR11LABEX0045DIGICOSME), operated by ANR as part of the program “Investissement d’Avenir” Idex ParisSaclay (ANR11IDEX000302).

