## Supplementary file for "State-of-the-RNArt: benchmarking current methods for RNA 3D structure prediction"

April 8, 2024

Table 1: Summary of the RNA molecules from Test Set I. It is obtained from RNAsolo [1] and Rfam [2].

| PDB ID | Rfam ID | Family | Sequence length |
|---|---|---|---|
| 1U9S | RF00010 | Gene; ribozyme | 155 |
| 1XJR | RF00164 | Cis-reg | 46 |
| 1YFG | RF00005 | tRNA | 64 |
| 2A64 | RF00011 | Gene;ribozyme | 298 |
| 2IL9 | RF00458 | Cis-reg | 135 |
| 2QUS | RF00008 | Gene;ribozyme | 68 |
| 3D2V | RF00059 | Cis-reg;riboswitch | 77 |
| 3DIL | RF00168 | Cis-reg;riboswitch | 173 |
| 3LA5 | RF00167 | Cis-reg;riboswitch | 71 |
| 3OXE | RF00504 | Cis-reg;riboswitch | 86 |
| 4AOB | RF00162 | Cis-reg;riboswitch | 94 |
| 4ENC | RF01734 | Cis-reg;riboswitch | 52 |
| 4LVW | RF01831 | Cis-reg;riboswitch | 89 |
| 4OQU | RF01725 | Cis-reg;riboswitch | 97 |
| 4PQV | RF01415 | Cis-reg | 68 |
| 4RUM | RF02683 | Cis-reg;riboswitch | 91 |
| 4WFL | RF01854 | Gene | 105 |
| 4ZNP | RF01750 | Cis-reg;riboswitch | 73 |
| 5NWQ | RF01763 | Cis-reg | 40 |
| 5T83 | RF00442 | Cis-reg;riboswitch | 89 |
| 6CU1 | RF02553 | Gene;sRNA | 79 |
| 6FZ0 | RF01826 | Cis-reg;riboswitch | 47 |
| 6N5P | RF00379 | Cis-reg;riboswitch | 126 |
| 6PRV | RF02541 | Gene;rRNA | 57 |
| 6UES | RF00634 | Cis-reg;riboswitch | 119 |
| 7D7W | RF03013 | Gene;sRNA | 51 |
| 7ELQ | RF03054 | Cis-reg | 45 |
| 7KGA | RF00525 | Cis-reg | 89 |
| 8SA3 | RF00174 | Cis-reg;riboswitch | 210 |

Table 2: Summary of the RNA molecules from Test Set II. It corresponds to single-stranded RNAs from RNA-Puzzles [3–7].

| **Puzzle name** | **PDB ID** | **Family** | **Sequence length** |
|---|---|---|---|
| rp03 | 3OWZ | Riboswitch | 84 |
| rp04 | 3V7E | Riboswitch | 126 |
| rp05 | 4P9R | Ribozyme | 188 |
| rp06 | 4GXY | Riboswitch | 168 |
| rp07 | 4R4V | Ribozyme | 185 |
| rp08 | 4L81 | Riboswitch | 96 |
| rp09 | 5KPY | Aptamer | 71 |
| rp11 | 5LYS | Riboregulator | 57 |
| rp12 | 4QLM | Riboswitch | 123 |
| rp13 | 4XW7 | Riboswitch | 71 |
| rp14_bound | 5DDO | Riboswitch | 61 |
| rp14_free | 5DDO | Riboswitch | 61 |
| rp16 | 6Y0Y | Ricin loop | 27 |
| rp17 | 5K7C | Ribozyme | 62 |
| rp18 | 5TPY | Virus | 71 |
| rp21 | 5NWQ | Riboswitch | 41 |
| rp23 | 6E8U | Aptamer | 37 |
| rp24 | 6OL3 | Virus | 112 |
| rp25 | 6P2H | Riboswitch | 69 |
| rp29 | 6TB7 | Riboswitch | 52 |
| rp32 | 7EOJ | Aptamer | 49 |
| rp34 | 7V9E | Ribozyme | 68 |

Table 3: Summary of the RNA molecules from Test Set III. It corresponds to the challenges from CASP-RNA [8].

| **Puzzle name** | **PDB ID** | **Type** | **Sequence length** |
|---|---|---|---|
| R1107 | 7QR4 | RNA | 69 |
| R1108 | 7QR3 | RNA | 69 |
| R1116 | 8S95 | RNA | 157 |
| R1117 | 8FZA | RNA/Ligand | 30 |
| R1126 | - | RNA | 363 |
| R1128 | 8BTZ | RNA | 238 |
| R1136 | 7ZJ4 | RNA | 374 |
| R1138 | 7PTK | RNA | 720 |
| R1149 | 8UYS | RNA | 124 |
| R1156 | 8UYE | RNA | 135 |
| R1189 | 7YR7 | RNA/protein | 173 |
| R1190 | 7YR6 | RNA/protein | 173 |

Table 4: Mean metrics for the benchmarked models for nine metrics for the Test Set I (RNAsolo). Decreasing metrics are separated by a double bar to increasing metrics.

| | **RMSD** | **MCQ** | **DI** | **P-VALUE** | **$\epsilon$RMSD** | **INF** | **TM-score** | **lDDT** | **GDT-TS** | **CAD-score** |
|---|---|---|---|---|---|---|---|---|---|---|
| MC-Sym | 15.27 | 36.56 | 34.83 | 0.11 | 2.05 | 0.44 | 0.24 | 0.01 | 0.22 | 0.27 |
| Vfold3D | 18.67 | 30.56 | 33.73 | 0.17 | 1.73 | 0.55 | 0.30 | 0.00 | 0.21 | 0.31 |
| RNAComposer | 21.78 | 25.04 | 36.46 | 0.24 | 1.63 | 0.60 | 0.25 | 0.43 | 0.20 | 0.32 |
| SimRNA | 23.00 | 25.50 | 38.89 | 0.29 | 1.83 | 0.59 | 0.25 | 0.03 | 0.18 | 0.31 |
| 3dRNA | 18.38 | 33.88 | 33.80 | 0.10 | 1.78 | 0.54 | 0.26 | 0.37 | 0.19 | 0.30 |
| IsRNA1 | 23.25 | **24.34** | 38.33 | 0.39 | 1.70 | 0.61 | 0.24 | 0.36 | 0.19 | 0.30 |
| RhoFold | 10.38 | 57.22 | 19.45 | **0.00** | 2.04 | 0.53 | 0.40 | 0.35 | 0.28 | 0.29 |
| trRosettaRNA | **5.21** | 32.75 | **9.32** | **0.00** | **1.47** | 0.56 | **0.65** | **0.63** | **0.44** | **0.39** |
| Vfold-Pipeline | 18.84 | 25.26 | 32.01 | 0.20 | 1.62 | 0.59 | 0.33 | 0.42 | 0.20 | 0.28 |
| RNAJP | 23.45 | 25.02 | 37.76 | 0.41 | 1.58 | **0.62** | 0.27 | 0.03 | 0.21 | 0.29 |

Table 5: Mean metrics for the benchmarked models for nine metrics for the Test Set II (RNA-Puzzles). Decreasing metrics are separated by a double bar to increasing metrics.

| | **RMSD** | **MCQ** | **DI** | **P-VALUE** | **$\epsilon$RMSD** | **INF** | **TM-score** | **lDDT** | **GDT-TS** | **CAD-score** |
|---|---|---|---|---|---|---|---|---|---|---|
| MC-Sym | 15.56 | 34.65 | 25.21 | 0.12 | 1.66 | 0.62 | 0.26 | 0.01 | 0.31 | 0.54 |
| Vfold3D | 18.02 | 30.30 | 32.00 | 0.15 | 1.77 | 0.56 | 0.25 | 0.04 | 0.28 | 0.53 |
| RNAComposer | 20.41 | 25.36 | 30.82 | 0.25 | 1.50 | 0.66 | 0.25 | 0.46 | 0.28 | 0.51 |
| SimRNA | 16.13 | **23.36** | 23.43 | **0.00** | 1.56 | **0.69** | 0.28 | 0.03 | 0.30 | 0.54 |
| 3dRNA | 14.24 | 31.45 | 21.74 | 0.03 | 1.56 | 0.65 | 0.31 | 0.47 | 0.32 | 0.53 |
| IsRNA1 | 18.76 | 24.39 | 27.78 | 0.24 | 1.52 | 0.68 | 0.25 | 0.43 | 0.28 | 0.52 |
| RhoFold | 9.06 | 52.90 | 14.30 | 0.01 | 1.47 | 0.63 | 0.41 | 0.51 | 0.40 | 0.57 |
| trRosettaRNA | **5.45** | 32.05 | **8.56** | **0.00** | **1.37** | 0.64 | **0.63** | **0.64** | **0.59** | **0.64** |
| Vfold-Pipeline | 16.78 | 23.82 | 24.15 | 0.19 | 1.42 | **0.69** | 0.36 | 0.47 | 0.37 | 0.58 |
| RNAJP | 22.37 | 25.85 | 37.82 | 0.19 | 1.62 | 0.59 | 0.23 | 0.06 | 0.25 | 0.48 |

Table 6: Mean metrics for the benchmarked models for nine metrics for the Test Set III (CASP-RNA). Decreasing metrics are separated by a double bar to increasing metrics.

| | **RMSD** | **MCQ** | **DI** | **P-VALUE** | **$\epsilon$RMSD** | **INF** | **TM-score** | **lDDT** | **GDT-TS** | **CAD-score** |
|---|---|---|---|---|---|---|---|---|---|---|
| Vfold3D | 22.03 | 33.24 | 41.05 | 0.11 | 1.58 | 0.54 | 0.30 | 0.01 | 0.23 | 0.61 |
| RNAComposer | 29.84 | 25.98 | 45.64 | 0.30 | 1.35 | 0.65 | 0.25 | **0.52** | 0.19 | 0.65 |
| SimRNA | 24.98 | **25.10** | 40.93 | 0.10 | 1.62 | 0.61 | 0.25 | 0.03 | 0.22 | 0.64 |
| 3dRNA | **16.28** | 33.82 | **27.61** | **0.00** | 1.50 | 0.59 | **0.29** | 0.46 | **0.27** | 0.61 |
| IsRNA1 | 21.58 | 25.66 | 34.57 | 0.08 | 1.46 | 0.62 | 0.25 | 0.42 | 0.24 | 0.62 |
| RhoFold | 19.20 | 61.80 | 44.62 | **0.00** | 2.26 | 0.43 | 0.25 | 0.20 | 0.24 | 0.56 |
| trRosettaRNA | 22.42 | 35.88 | 43.14 | 0.01 | 1.59 | 0.52 | 0.27 | 0.49 | 0.23 | **0.67** |
| Vfold-Pipeline | 22.92 | 29.82 | 40.25 | 0.24 | 1.46 | 0.57 | 0.28 | 0.47 | 0.24 | 0.60 |
| RNAJP | 23.55 | 25.30 | 34.42 | 0.23 | **1.26** | **0.68** | 0.28 | 0.04 | 0.26 | 0.63 |

Table 7: Mean metrics for the benchmarked models for nine metrics for the pooled test sets. Decreasing metrics are separated by a double bar to increasing metrics.

| | **RMSD** | **MCQ** | **DI** | **P-VALUE** | **$\epsilon$RMSD** | **INF** | **TM-score** | **lDDT** | **GDT-TS** | **CAD-score** |
|---|---|---|---|---|---|---|---|---|---|---|
| MC-Sym | 15.52 | 35.38 | 28.67 | 0.14 | 1.82 | 0.54 | 0.25 | 0.01 | 0.28 | 0.44 |
| Vfold3D | 18.88 | 30.78 | 34.04 | 0.15 | 1.72 | 0.55 | 0.28 | 0.02 | 0.24 | 0.44 |
| RNAComposer | 21.48 | 25.46 | 34.48 | 0.27 | 1.55 | 0.62 | 0.25 | 0.45 | 0.23 | 0.43 |
| SimRNA | 20.85 | 24.94 | 33.76 | 0.18 | 1.73 | 0.62 | 0.26 | 0.03 | 0.22 | 0.44 |
| 3dRNA | 16.45 | 32.72 | 27.67 | 0.06 | 1.66 | 0.59 | 0.29 | 0.41 | 0.25 | 0.44 |
| IsRNA1 | 21.22 | **24.52** | 33.41 | 0.29 | 1.60 | **0.64** | 0.25 | 0.39 | 0.23 | 0.43 |
| RhoFold | 11.08 | 56.15 | 19.91 | **0.00** | 1.85 | 0.56 | 0.39 | 0.40 | 0.32 | 0.43 |
| trRosettaRNA | **6.79** | 33.08 | **11.79** | **0.00** | **1.45** | 0.58 | **0.59** | **0.61** | **0.47** | **0.51** |
| Vfold-Pipeline | 18.57 | 25.20 | 29.67 | 0.20 | 1.53 | 0.63 | 0.33 | 0.45 | 0.27 | 0.44 |
| RNAJP | 23.04 | 25.38 | 37.33 | 0.30 | 1.56 | 0.62 | 0.26 | 0.04 | 0.23 | 0.41 |

| | **RMSD** | **MCQ** | **$\epsilon$RMSD** | **DI** | **P-VALUE** | **TM-SCORE** | **GDT- TS** | **INF** | **lDDT** | **CAD-score** |
|---|---|---|---|---|---|---|---|---|---|---|
| MC-Sym [9] | 15.96 | 34.84 | 1.38 | 21.47 | 0.00 | 0.27 | 0.32 | 0.74 | 0.02 | 0.67 |
| Vfold3D [10] | 15.79 | 30.87 | 1.56 | 24.83 | 0.00 | 0.30 | 0.35 | 0.64 | 0.01 | 0.64 |
| RNAComposer [11] | 18.27 | 19.38 | 1.20 | 23.69 | 0.00 | 0.33 | 0.36 | 0.77 | 0.55 | 0.74 |
| SimRNA [12] | 17.7 | 23.79 | 1.58 | 25.16 | 0.00 | 0.30 | 0.30 | 0.70 | 0.07 | 0.70 |
| 3dRNA [13] | 9.56 | 33.28 | 1.37 | 13.47 | 0.00 | 0.41 | 0.42 | 0.71 | 0.50 | 0.70 |
| IsRNA1 [14] | 11.75 | 23.74 | 1.40 | 16.37 | 0.00 | 0.29 | 0.34 | 0.72 | 0.45 | 0.70 |
| RhoFold [15] | 4.10 | 50.86 | 1.20 | 5.83 | 0.00 | 0.60 | 0.59 | 0.70 | 0.56 | 0.76 |
| trRosettaRNA [16] | **2.38** | 24.91 | **0.90** | **2.85** | 0.00 | **0.80** | **0.82** | 0.83 | **0.73** | **0.86** |
| Vfold-Pipeline [17] | 6.52 | 18.3 | 0.99 | 7.63 | 0.00 | 0.49 | 0.55 | **0.85** | 0.69 | 0.81 |
| RNAJP [18] | 22.48 | 36.77 | 1.69 | 40.89 | 0.01 | 0.23 | 0.26 | 0.55 | 0.01 | 0.58 |

Table 8: Metrics for the ten benchmarked models for RNA challenge 3 (rp03) (id: 3OWZ, 84 nucleotides). Decreasing metrics are separated by a double bar to increasing metrics.

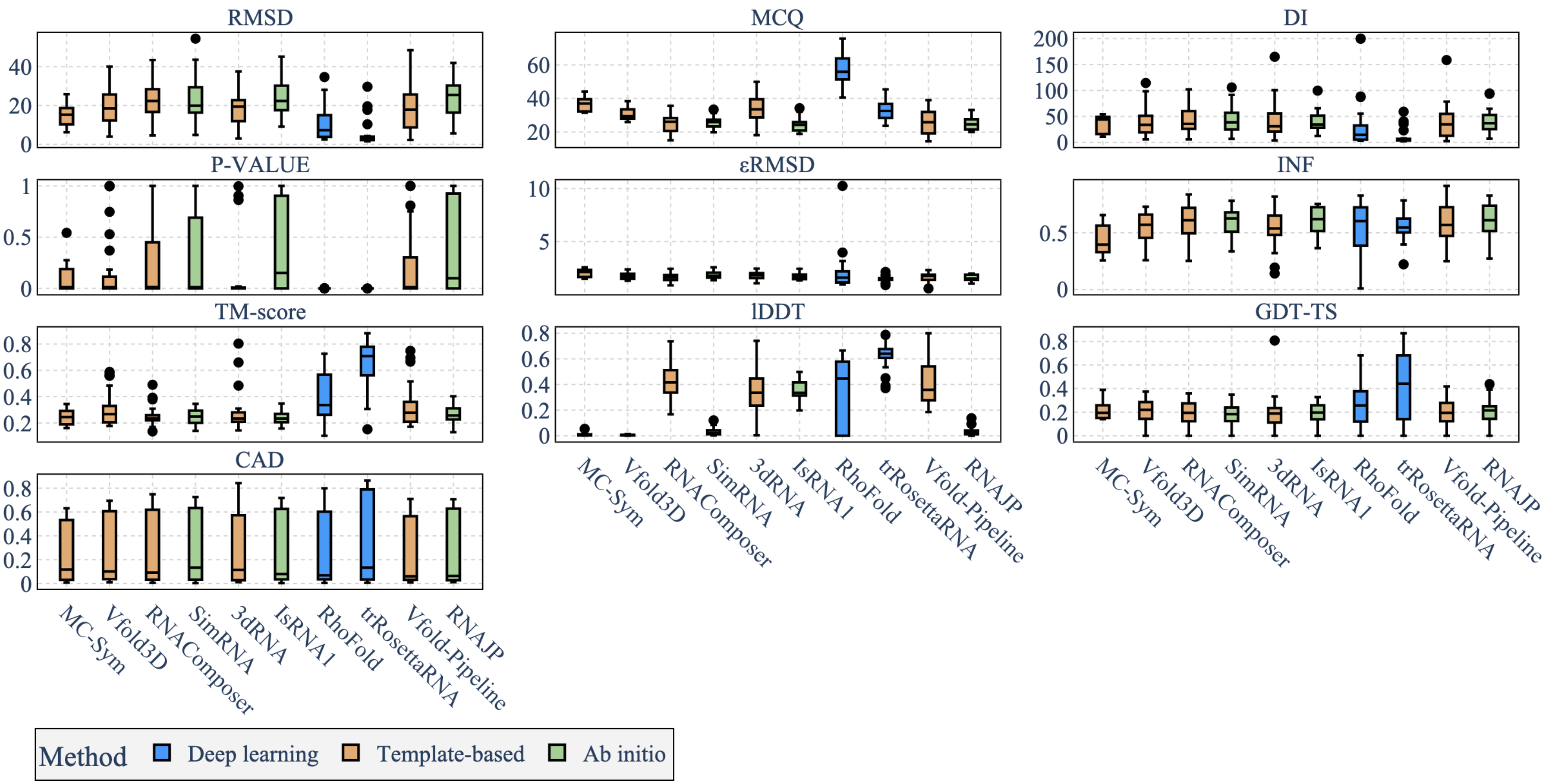


Figure 1: Distribution of the different metrics on Test Set I (RNAsolo) for the ten benchmarked models. RMSD, MCQ, DI, P-VALUE, and $\epsilon$RMSD are decreasing metrics, whereas INF, TM-score, GDT-TS, CAD-score and lDDT are increasing (the higher, the better).

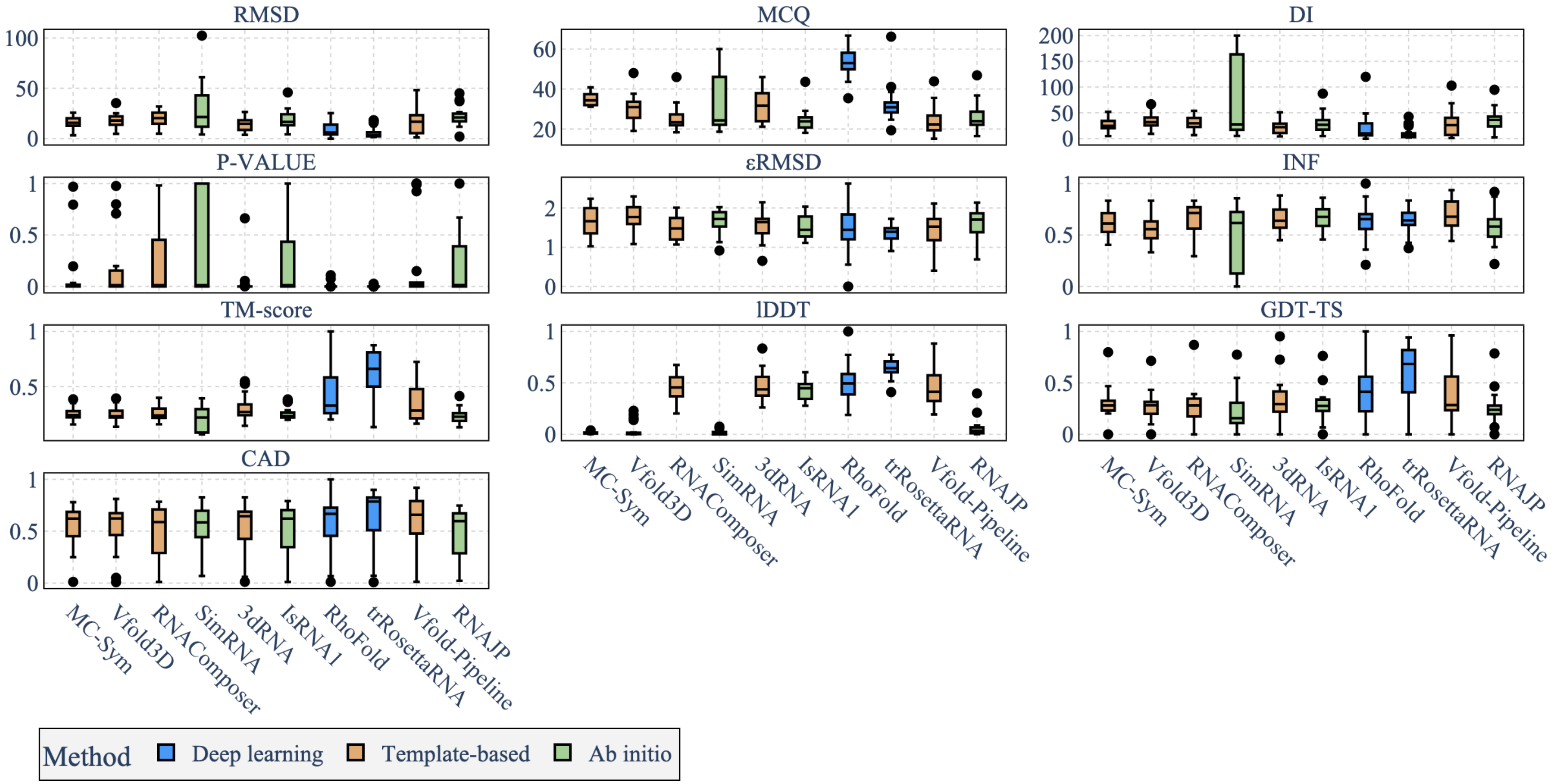


Figure 2: Distribution of the different metrics on Test Set II (RNA-Puzzles) for the ten benchmarked models. RMSD, MCQ, DI, P-VALUE, and $\epsilon$RMSD are decreasing metrics, whereas INF, TM-score, GDT-TS, CAD-score and lDDT are increasing (the higher, the better).

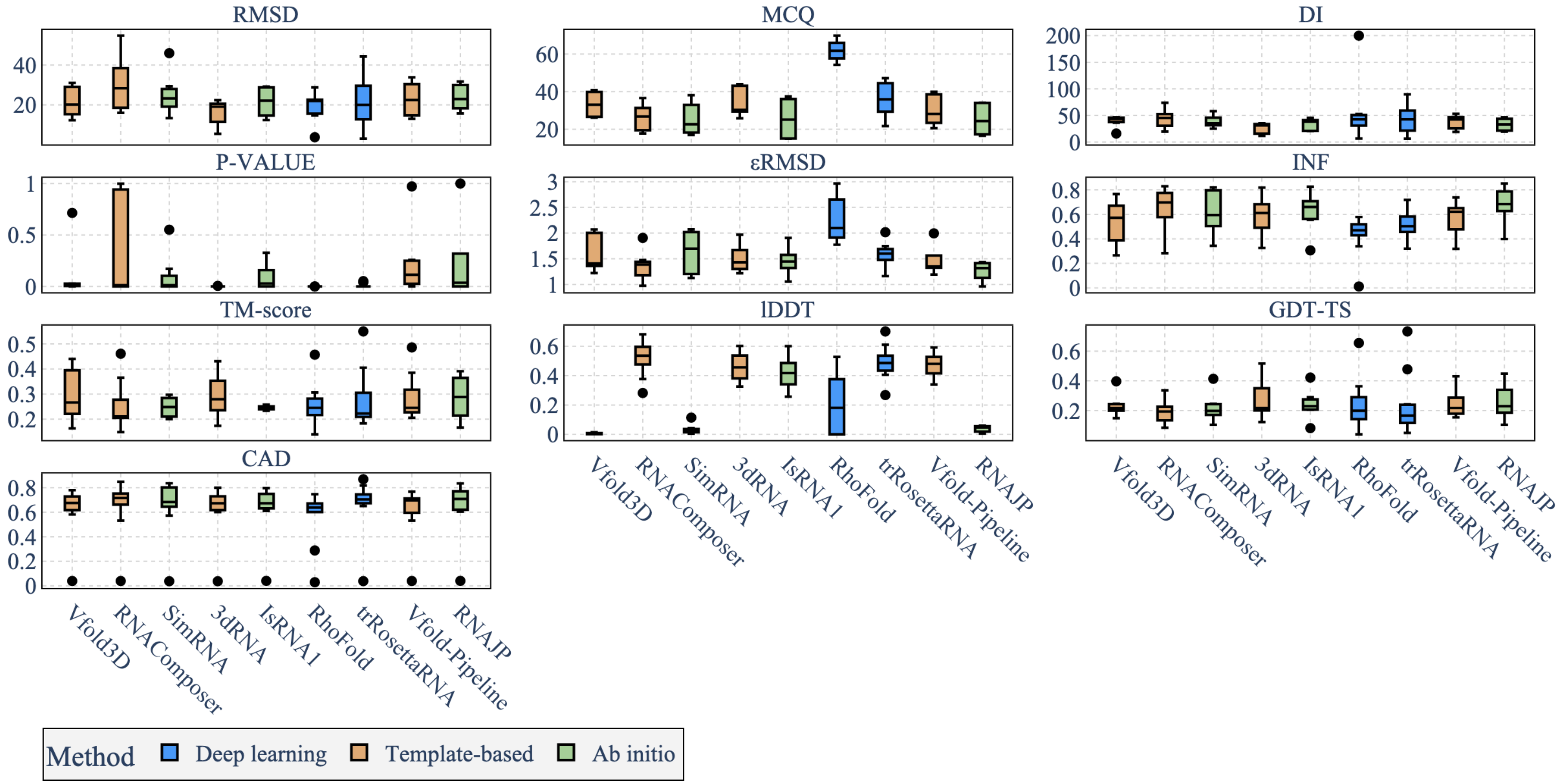


Figure 3: Distribution of the different metrics on Test Set III (CASP-RNA) for the ten benchmarked models. RMSD, MCQ, DI, P-VALUE, and $\epsilon$RMSD are decreasing metrics, whereas INF, TM-score, GDT-TS, CAD-score and lDDT are increasing (the higher, the better).

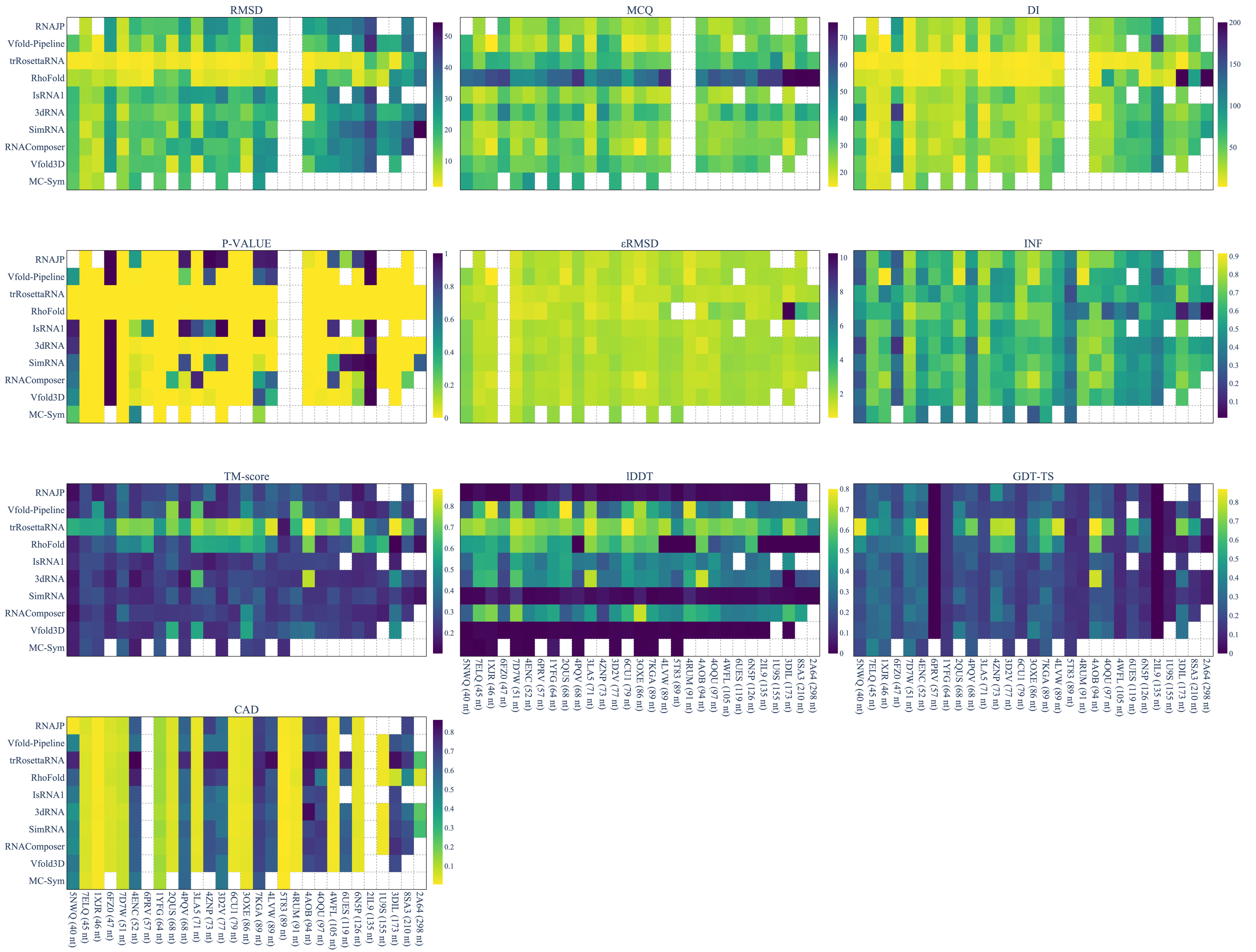


Figure 4: Prediction results obtained by the ten benchmarked methods on each RNA of the Test Set I (RNAsolo) dataset. The results are reported using different metrics: RMSD, MCQ, DI, P-VALUE, $\epsilon$RMSD, INF, TM-score, lDDT, CAD-score and GDT-TS. The best results are in yellow, while the bad results are in dark. Missing values (in white) are due to a failure in predictions by the models or in the metric computation. Challenges are sorted by RNA sequence length. RNA length is provided in brackets.

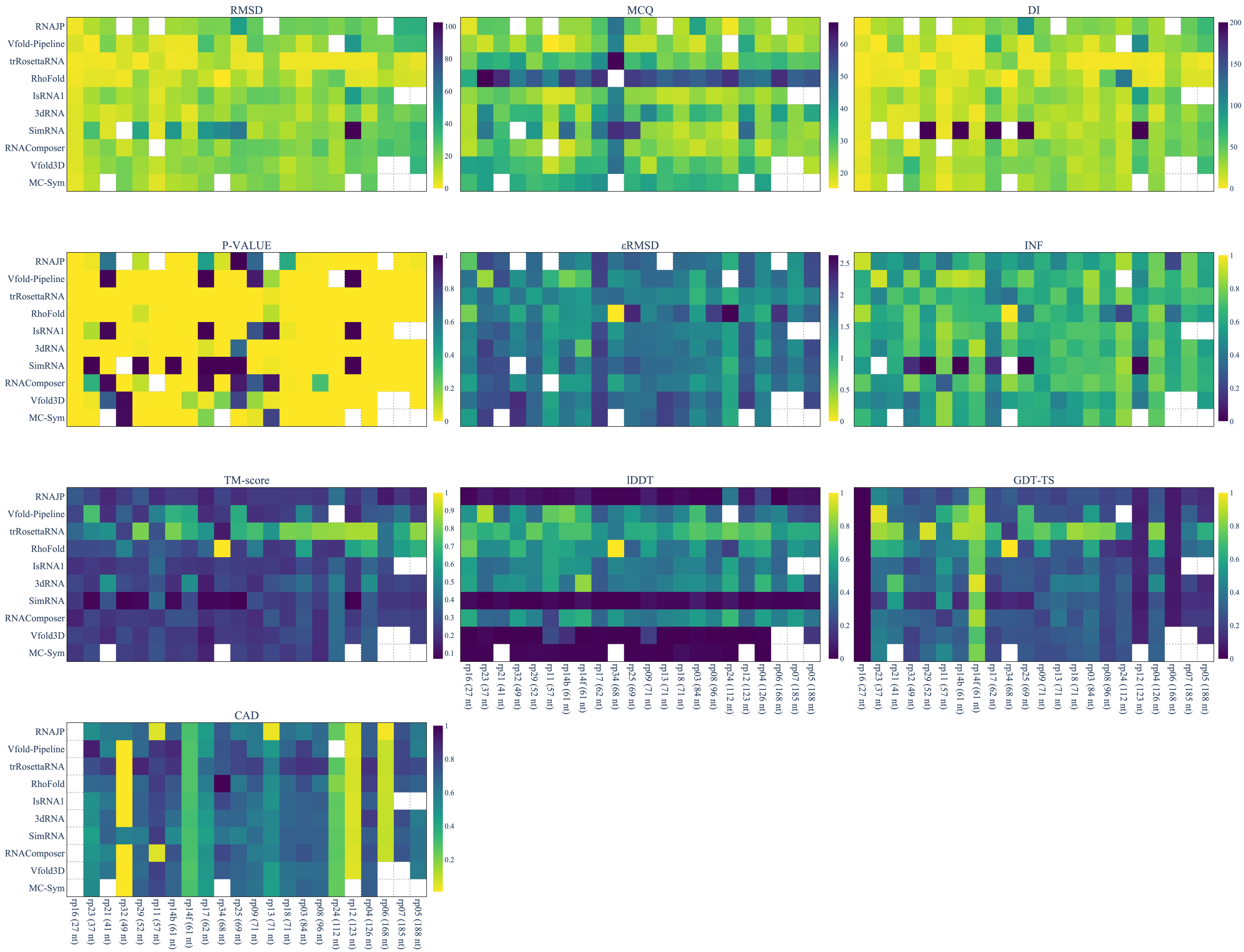


Figure 5: Prediction results obtained by the ten benchmarked methods on each RNA of the Test Set II (RNA-Puzzles) dataset. The results are reported using different metrics: RMSD, MCQ, DI, P-VALUE, ϵRMSD, INF, TM-score, lDDT, CAD-score and GDT-TS. The best results are in yellow, while the bad results are in dark. Missing values (in white) are due to a failure in predictions by the models or in the metric computation. Challenges are sorted by RNA sequence length. RNA length is provided in brackets.

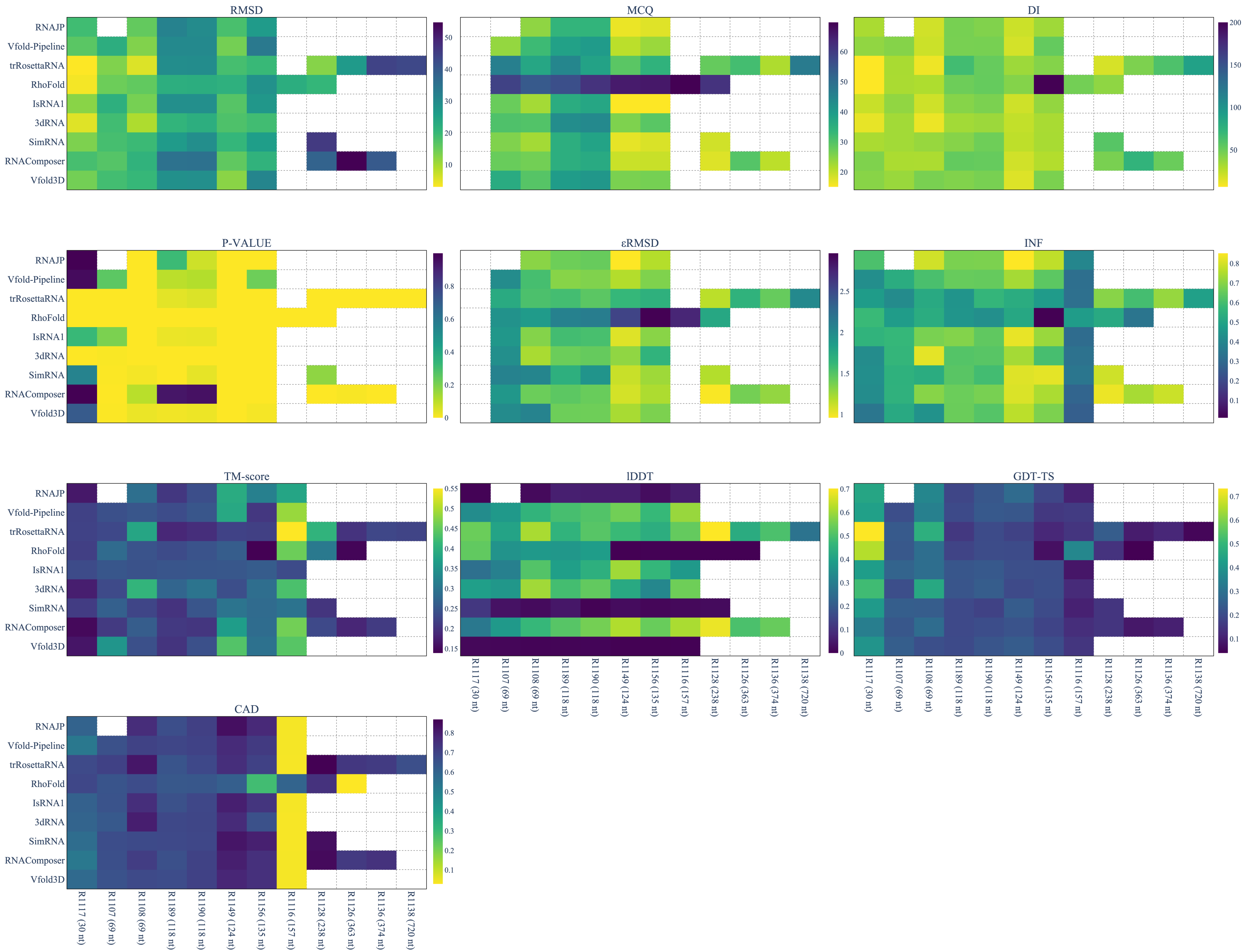


Figure 6: Prediction results obtained by the ten benchmarked methods on each RNA of the Test Set III (CASP-RNA) dataset. The results are reported using different metrics: RMSD, MCQ, DI, P-VALUE, ϵRMSD, INF, TM-score, lDDT, CAD-score and GDT-TS. The best results are in yellow, while the bad results are in dark. Missing values (in white) are due to a failure in predictions by the models or in the metric computation. Challenges are sorted by RNA sequence length. RNA length is provided in brackets.
